# Evaluation of eight protocols for genomic DNA extraction of *Hypostomus commersoni* Valenciennes, 1836 (Loricariidae: Siluriformes)

**DOI:** 10.1101/744946

**Authors:** P. Mezzomo, A. A. Mielniczki-Pereira, T. L. Sausen, J. R. Marinho, R. L. Cansian

## Abstract

The principle and the techniques applied in DNA extraction play a pivotal role in the obtention of a purified genetic material. The present study investigates the efficiency of eight protocols in the DNA extraction of *Hypostomus commersoni*, an essential component of South American freshwater ichthyofauna. The quality of samples was assessed through spectrophotometry, gel electrophoresis, and PCR-RAPD amplification. The efficiency of DNA extraction was influenced both by the method applied and the target-tissue of choice. Higher concentrations and yield of DNA were obtained from ocular tissue, with a positive spectrum of incubation in lysis buffer for up to 36 hours after sample collection, using fresh tissues and in the presence of a high concentration of Proteinase K (20 mg.ml^-1^). In these conditions, samples were successfully amplified. To date, there is no record of description for the parameters analyzed in this work, neither the description of RAPD markers for the species *H. commersoni*.

## 1. Introduction

A crucial step for high-quality genetic analysis is considered to be a satisfactory DNA extraction. Nevertheless, this purpose is not always easy to achieve, since different biological compounds, such as lipids and proteins, can act as contaminants and interfere in the final quality of the product (Bauer and Patzelt, 2003). The principles and techniques applied in DNA extraction play a pivotal role in obtaining a considerable and purified yield of this molecule (Muhammad et al., 2016).

A variety of methods has been established to isolate DNA from biological materials (Milligan, 1998; Reischl et al., 2000; Rantakokko-Jalava, 2002; Tongeren et al., 2011). Regardless of the method of choice, the procedure usually consists of three steps: lysis, purification, and DNA recovery; being the first stage the most critical one since it is at this point that the chances of causing DNA damages are higher (Barbosa et al., 2015).

The quality and amount of available DNA strand template have a significant influence on genetic analyzes based on PCR (Polymerase Chain Reaction), for example (Ward et al., 2009; Wang et al., 2011). Therefore, an ideal extraction technique should optimize DNA yield, minimize DNA degradation, and be efficient in terms of cost, time, labor, and supplies (Chen et al., 2010).

With the availability of a plethora of molecular analysis machinery, there is an exponential demand for high-quality samples, capable of generating great results with a minimum of errors. To accomplish this, commercial kits were developed, aiming for DNA extractions of excellence (Coombs et al., 1999; Barea et al., 2004; Bueno, 2004).

However, the market price of these products is not always easily affordable (Chen et al., 2010; Mega, Revers, 2011). On the other hand, the extraction of genetic material using reagents of homemade preparation can be less costly, and, therefore, such methodologies can be replicated in laboratories with a limited budget.

Establish a well-succeed protocol for DNA extraction is a crucial step in genetic analysis, considering that the same methodology may present variations between different organisms (Lucentini et al., 2006). Furthermore, the quality of isolated DNA through methods described for other taxa generally presents unsatisfactory results when applied to fish species (Muhammad et al., 2016).

*Hypostomus commersoni* (Valenciennes, 1836), commonly known as suckermouth catfish, is an essential component of freshwater ichthyofauna, inhabiting the Paraná and Uruguay basins, in South America (Burgess, 1989; Reis et al., 2003). This taxon is marked by its ecological and commercial relevance (Buck and Sazima, 1995; Garavello and Garavello, 2004; Baldisserotto, 2009; Gonçalves, 2011; Lujan et al., 2012), and, despite that, there is no record of description of well-succeed DNA extraction protocol nor the description of PCR-RAPD markers for this organism to date. In this work, we evaluated eight protocols in the extraction of the genomic DNA of *H. commersoni*, aiming to obtain high-quality DNA samples of this specie.

## 2. Material and methods

### 2.1 Ethics Statement

This study was approved by the CEUA (Ethics Committee for Animal Use, Protocol nº 79) of URI (Universidade Regional Integrada). Written informed consent was obtained from the regulatory organ.

### 2.2 Sample collection

Forty specimens of *Hypostomus commersoni* (24 cm ± 114 g) were collected in a small tributary from the Upper Rio Uruguay basin in the municipality of Erechim, South Brazil. The fishes were subjected to euthanasia, using clove oil in a concentration of 0,1%. Specimens were allocated in a recipient containing local river water, in which the clove oil was applied. The tissues previously selected for DNA extraction were removed and placed in a sterile 2 mL Eppendorf tube for subsequent analysis. Vouchers specimens were deposited in the ichthyological collection of the Museu Regional do Alto Uruguai (MuRAU) of URI-Erechim, Brazil (MuRAU - PEIXES 715-716).

### 2.3 DNA extraction

All DNA was extracted and quantified at the biochemistry and molecular biology laboratory of URI. Seven protocols for DNA extraction were selected from the literature, whose methodologies had previously been described for fish species or whose parameters were similar to those intended to be analyzed in this work, such as the same extraction conditions or target-tissues (Doyle and Doyle, 1987; Medrano et al., 1990; Whitmore et al., 1992; Aljanabi and Martinez, 1997; Faleiro et al., 2003; Barrero et al., 2008; Mossi et al., 2014). An eighth protocol was proposed and evaluated, combining the variables that presented better results in the extraction of *H. commersoni* DNA in each of the seven tested protocols. The full description of the protocols one to seven is available as supplementary material (S1). The eighth protocol is described in full in the results section.

In combination with the protocols, we also evaluate eight different variables that, based on literature findings and our previous analysis, were thought to influence the purity and amount of the obtained DNA, as follows:

a. **Forms of sample preservation:**
  1. *Fresh tissues*: When the specimen is collected, immediately submitted to euthanasia, and right after the tissue is removed and submitted to DNA extraction.
  2. *Tissues stored in 70% and 95% ethanol:* When the specimen is collected, immediately submitted to euthanasia, and right after the tissue is removed, and stored in tubes containing ethanol until its use, at room temperature (approximately 27 ºC).
  3. *Tissues stored at −10 °C or −20 °C*: When the specimen is collected, immediately submitted to euthanasia, and tissues are removed and directly frozen until its use.
b. **Forms of sample maceration:**
  1. *Chemical lysis:* Samples submerse in lysis buffer, tubes placed on a water bath at 27ºC, without mixing.
  2. *Mechanical lysis:* Samples submersed in lysis buffer and mixed using the vortex.
  3. *Lysis with liquid nitrogen (LN2):* Samples triturated; fragments placed in tubes with lysis buffer on a water bath at 27ºC.
  4. *Manual lysis:* Samples smashed using a sterile glass stick, and then placed in tubes with lysis buffer on a water bath at 27 ºC.
c. **Different target-tissues**: dorsal fin, caudal fin, muscle, gills, blood, and ocular tissue.
d. **Times of incubation in extraction buffer**: one hour (55 ºC), 12 hours (27 ºC), 24 hours (27 ºC), and 36 hours (27 ºC), for all the conditions mentioned in items a and b.
e. **RNAse:** presence or absence.
f. **Proteinase K:** presence or absence, with gradients of concentration (05 mg.ml-1, 1 mg.ml-1, 20 mg.ml-1 and 200 mg.ml-1).
g. **Storage after extraction**: at 4 ºC, −10 ºC or −20 ºC.
h. **Integrity and quantification:** on the day of extraction, after 10, 20, and 30 days of storage.

Several tests aiming for experimental optimization were performed, and for each variable, the factors that implicated in better concentration and purity of DNA were selected. The results were categorized into a qualitative matrix, according to the effect on the final extracted product: a. Efficient: satisfactory concentration and purity; b. Indifferent: did not influence the parameters evaluated; c. Inefficient: low concentration of genetic material, low purity, or degradation of samples.

### 2.4 Analysis of DNA integrity

The integrity of the samples was verified through agarose gel (0,8%) electrophoresis, stained with ethidium bromide (1 mg/ml). The reaction runs for 150 minutes at 90 volts, and the visualization and image caption of the gel was obtained under ultraviolet (UV) light.

### 2.5 Evaluation of concentration and purity of DNA

The samples whose extraction was successful according to the electrophoresis verification were then submitted to spectrophotometry and analyzed for protein contamination (purity). Samples were quantified in a spectrophotometer (model NI 1600UV - Nova Instruments) at the wavelengths of A_260_ and A_280_ nanometers (nm). The purity of the DNA was confirmed by the ratio of absorbance values at the two wavelengths measured (Barbosa, 1998). The DNA concentration of each sample was determined by the equation: [DNA] = 50 µg/ml x D (dilution factor) x A_260_ (read value obtained in the wavelength of 260 nm). Satisfactory purity values were considered to be ranging between 1,7-2,0.

### 2.6 DNA amplification by PCR-RAPD

The final quality assessment of the samples was verified through their amplification with RAPD markers. Samples that reached the above parameters of DNA concentration and yield were submitted to amplification with different primers of PCR-RAPD. In total, 32 primers from Operon Technologies sets OPA (01, 02, 03, 04, 05, 06, 07, 08, 09), OPB (03, 05, 07, 10, 11, 15), OPF (10, 11, 12 e 13), OPW (01, 02, 03, 04, 05, 06, 07, 08, 09, 10) and OPY (01,02,03) were tested. Since the RAPD technique is sensitive to changes in reaction parameters (e.g., primer, MgCl2, dNTP concentrations) (Chiappero and Gardenal, 2001), the same reaction conditions were used for all samples. Ten nanograms of genomic DNA was amplified in PCR buffer (20 mM Tris-HCl pH 8,4, 50 mM KCl), 4 mM MgCl_2_, 0.7 mM dNTPs, 0,3 μM primers e one unity of Taq DNA polymerase enzyme (Invitrogen). The reaction was conducted in a final volume of 25 µL, in the following conditions: denaturation at 95ºC for 30 seconds, annealing at 50 ºC for 1 min, and extension at 72 ºC for 5 min totaling 40 cycles. The amplification products were applied in agarose gel (1,5%), stained with ethidium bromide (1 mg/ml). The reaction runs for 150 minutes at 150V. Visualization and image caption of band patterns were obtained under ultraviolet (UV) light.

*2.7 Data analysis*

Statistical analysis was performed using the Software Python (version 3.6.0). Differences between DNA extraction methods in total DNA concentration were determined by ANOVA using Tukey HSD as a post-hoc test.

## 3. Results

### 3.1 DNA extraction method impacts DNA yield

Our study has shown that a method that uses in-house produced reagents instead of commercial kits for DNA extraction can generate samples with sufficient quality to be used in molecular analysis, as PCR-RAPD. The amount of DNA extracted was dependent on the technique and the target-tissue of choice. For all variables examined, the improved extraction method developed and proposed at this work, produced the highest total DNA mean yields, for all tissues, resulting in significantly higher absorbance ratios when compared to the other extraction methods evaluated. The influence of each variable evaluated on the final DNA extraction is available in Table 1.

**Table 1.**
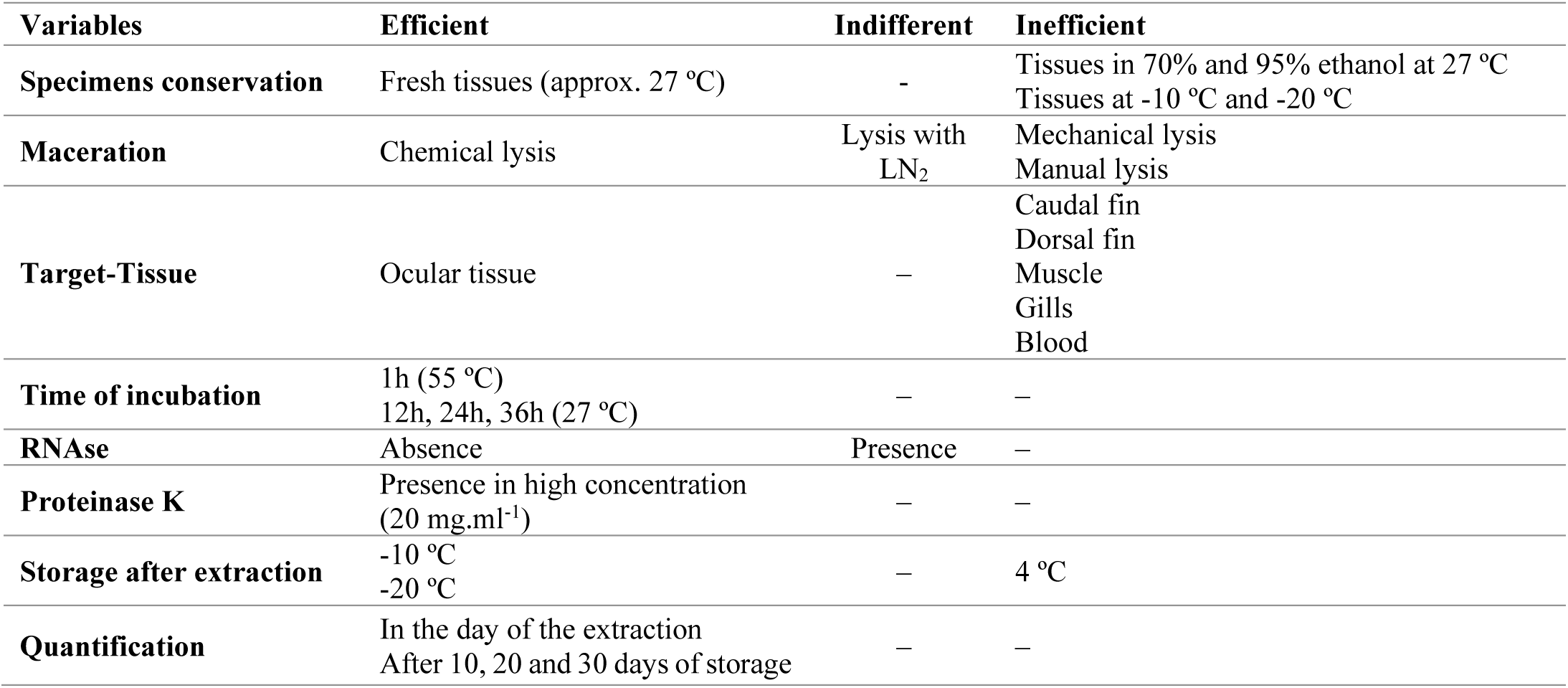
Effect of different variables in the total yield of extracted DNA from *H. commersoni*.

### 3.2 Improved protocol for DNA extraction of Hypostomus commersoni

The protocol proposed in this work presented, in general, the highest efficiency in the extraction of *H. commersoni* DNA. The samples submitted to extraction through this protocol were the only ones that amplified with PCR-RAPD primers. The complete methodology of this protocol is described in Table 2.

**Table 2.** Description of reagents and steps for DNA extraction of *H. commersoni* through the method proposed in this work.

| Reagents |  |
| --- | --- |
| STE Buffer | 0,1 M NaCl + 10 mM Tris + 1 mM EDTA (pH 8,0) |
| Lysis Buffer | 2,5 mM Tris-HCl pH 8,0 + 0,1 mM EDTA pH 8,0 + 0,1% SDS + 8,0 mM NaCl |
| Proteinase K | 20 mg.mL<sup>-1</sup> |
| CIA | Chloroform + isoamyl alcohol (1:24 v/v) |
| TE Buffer | 10 mM Tris pH 8,0 + 1 mM EDTA pH 8,0 |
| TBE Buffer | 500 mM Tris-HCl + 60 mM boric acid + 83 mM EDTA |

### 3.3 Analysis of DNA integrity in an electrophoresis gel

All samples obtained from the six tissues and tested under the different conditions were submitted to gel electrophoresis in triplicate, resulting in a total count of 144 samples. The samples obtained from ocular tissue presented the highest integrity of extracted DNA, in all combinations of variables and protocols. For all tissues, the leading bands of DNA were around 46-48 kb in size. Comparing protocols, the EDTA methods (protocols two, seven, and eight) showed relatively lighter smear tails, indicating less DNA degradation.

### 3.4 Quantification and assessment of purity

The resulting values of quantification for each DNA sample extracted under the different conditions and through the different protocols are presented in Tables 3 and 4, as well as in Figures 1A–1F. The amount of DNA obtained, and final yield mean values varied between samples from different tissues, as well as between samples of the same tissue but extracted using different protocols.

**Table 3.**
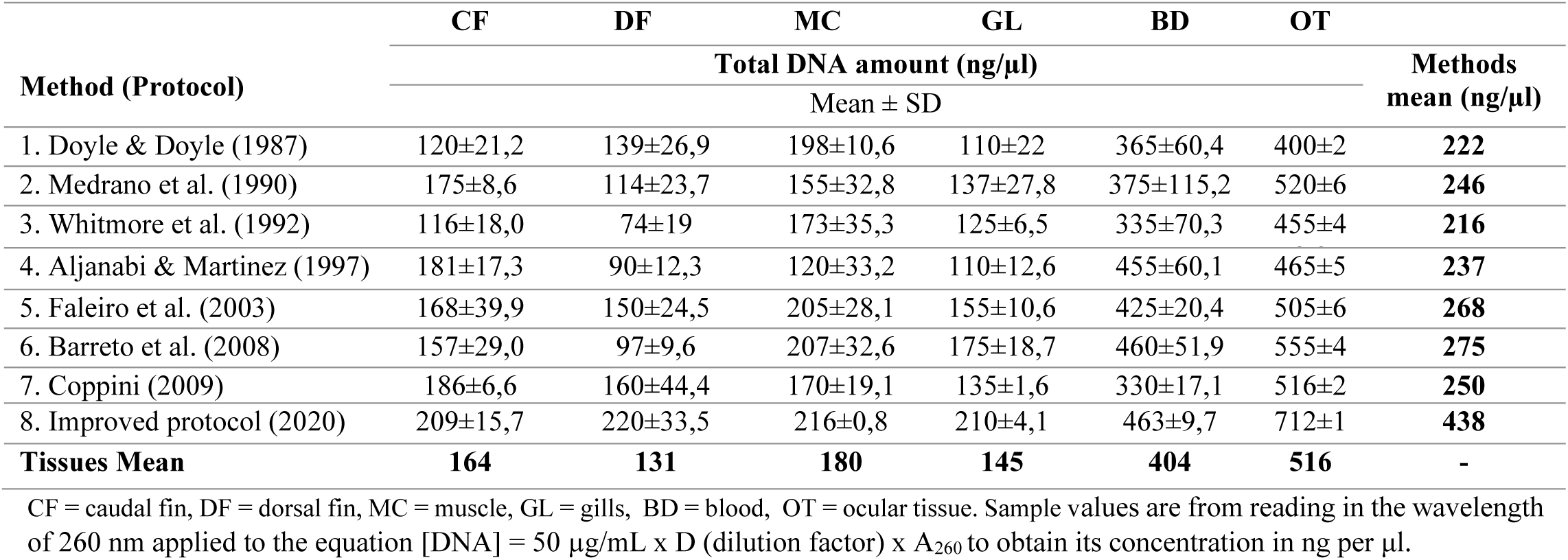
Mean of total DNA amount (ng/μl) extracted using eight different methods for six different tissues.

**Table 4.** The mean of DNA amount from ratio 260/280 nm, extracted using eight different methods for six different tissues.

| Method (Protocol) | CF | DF | MC | GL | BD | OT | Methods<br>Mean |
| --- | --- | --- | --- | --- | --- | --- | --- |
|  | Absorbance ratio (OD 280/260 nm) |  |  |  |  |  |  |
|  | Mean ± SD |  |  |  |  |  |  |
| 1, Doyle & Doyle (1987) | 1,5 ±0,016 | 0,8 ±0,08 | 1,4 ±0,08 | 1 ±0 | 1,9 ±0,08 | 1,8 ±0,12 | **1,4** |
| 2, Medrano et al. (1990) | 1,8 ±0,08 | 0,6 ±0,08 | 1,6 ±0,08 | 1 ±0,08 | 1,4 ±0,08 | 1,6 ±0,08 | **1,3** |
| 3, Whitmore et al. (1992) | 1 ±0,14 | 0,7 ±0,08 | 1,6 ±0,08 | 1,2 ±0 | 1,6 ±0,08 | 1,8 ±0,08 | **1,3** |
| 4, Aljanabi & Martinez (1997) | 1,1 ±0,16 | 0,8 ±0,08 | 1 ±0,08 | 1 ±0 | 1,7 ±0,08 | 1,6 ±0,08 | **1,2** |
| 5, Faleiro et al. (2003) | 1,1 ±0,08 | 0,9 ±0,08 | 1,6 ±0,08 | 1,1 ±0,14 | 1,6 ±0 | 1,9 ±0,08 | **1,4** |
| 6, Barreto et al. (2008) | 1,1 ±0,08 | 0,9 ±0,08 | 1,3 ±0,08 | 1,2 ±0 | 1,8 ±0,08 | 1,7 ±0,08 | **1,3** |
| 7, Coppini (2009) | 1,5 ±0,08 | 1,1 ±0,12 | 0,8 ±0,08 | 1,1 ±0,08 | 1,6 ±0 | 1,8 ±0,08 | **1,3** |
| 8, Improved protocol (2019) | 2 ±0 | 1,5 ±0,08 | 1,7 ±0,08 | 1,5 ±0,08 | 1,9 ±0,08 | 2 ±0,08 | **1,8** |
| **Tissues Mean** | **1.4** | **0.9** | **1.3** | **1.1** | **1.7** | **1.8** | **-** |

CF = caudal fin, DF = dorsal fin, MC = muscle, GL = gills, BD = blood, OT = ocular tissue, Sample values are from the ratio of the reads in the wavelength of 260 and 280 nanometers, that quantifies the purity of DNA in the samples.

**Figure 1a-f.**
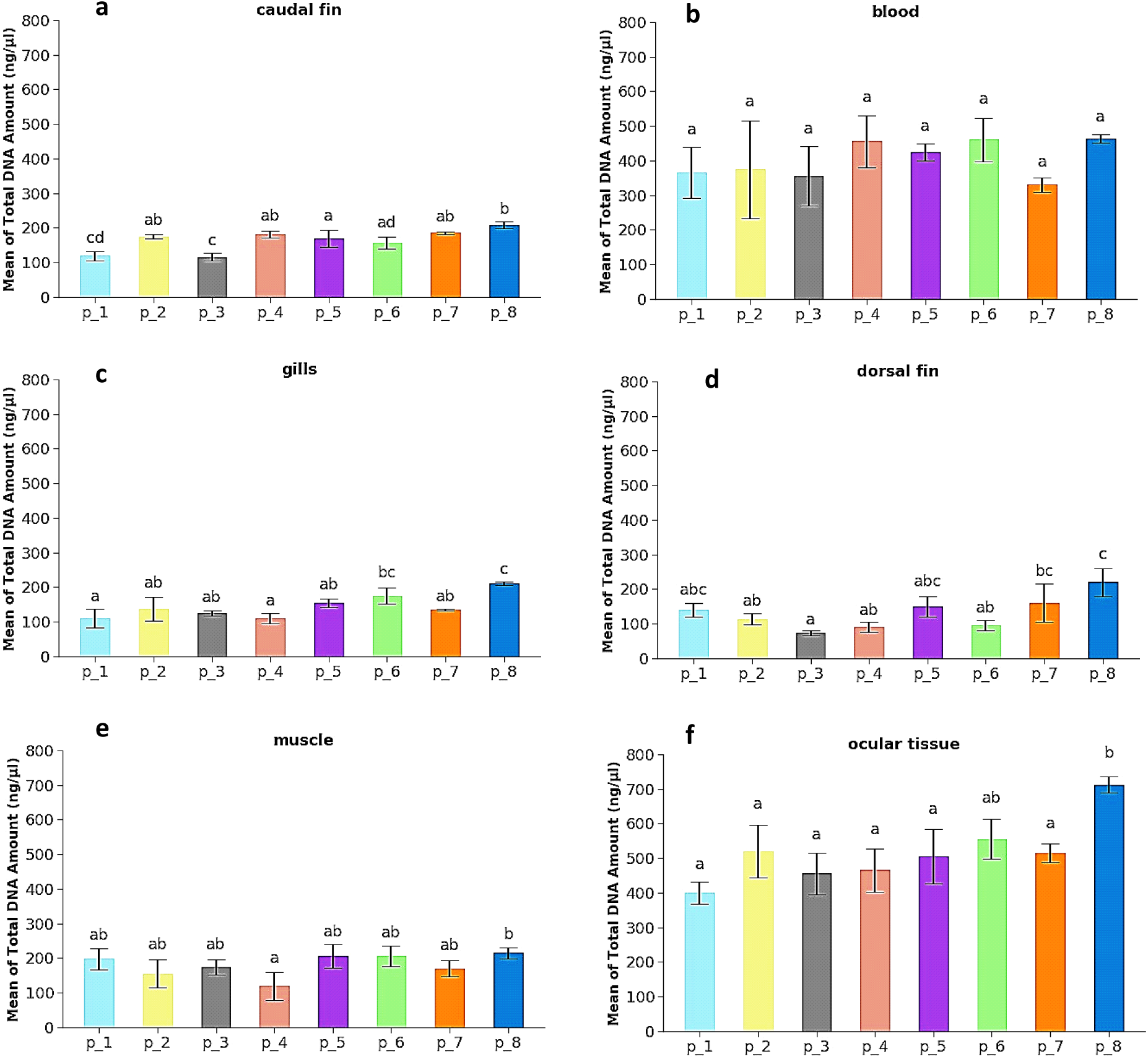
Genomic DNA extraction efficiency of *Hypostomus commersoni* varies dependent on the extraction method and target tissue. Mean total amount (± standard deviation) of DNA extracted from caudal fin (a), blood (b), gills (c), dorsal fin (d), muscle (e), and ocular tissue (f), using eight DNA extraction protocols. Samples were extracted, and DNA was quantified by spectrometry. Statistical significance was determined using one way ANOVA with Tukey HSD post hoc test. Significance defined by p ≤ 0.05 and denoted by lower case letters. Whiskers denoting lower standard deviation are cropped at zero.

When comparing samples, the ocular tissue had the highest mean yield of extracted DNA (516 ng/μl), while the lowest concentration was for the dorsal fin (131 ng/μl). Concerning the protocols, means between different tissues indicated that the lowest value was for extractions performed using protocol three (216 ng/μl), while the highest mean concentration of DNA was obtained with protocol eight (438 ng/μl). Statistically, there is no significant difference between most of the samples whose DNA was extracted from the tissue’s dorsal fin, caudal fin, gills, and muscle. The major exceptions were blood and ocular tissue, which had similar p-values for most of the evaluated protocols.

Although the results were statistically similar when concerning the total amount of DNA obtained per sample, the mean values of the absorbance ratio presented marked variance. The values for the absorbance ratio were 1,2 (protocol four), 1,3 (protocols two, three, six and seven), 1,4 (protocols one and five), and 1,8 (protocol eight), indicating its higher efficiency over the other seven protocols.

### 3.5 DNA extraction method influences PCR-RAPD amplification

RAPD markers have been used to evaluate the genetic diversity in numerous organisms (Cooper, 2000; Ali et al., 2004; Torezan et al., 2005; Bickel et al., 2006). Studies on genetic conservation of fish populations from South American rivers have successfully applied such markers to access the genetic diversity of different fish species (Almeida et al., 2001; Dergam et al., 2002; Wasko and Galetti Jr, 2002; Leuzzi et al., 2004; Matoso et al., 2004; Wasko et al., 2004). Of the thirty-four primers tested, nine amplified the DNA of the evaluated species in the concentrations of 40 and 20 ng/µl. These can be applied in studies evaluating the genetic structure of *H. commersoni*. The score of bands varied between the amplification using different concentrations of DNA (Table 5).

**Table 5.** Score of DNA bands amplified for each primer.

| Primer | Sequence 5' - 3' | Score of Amplified DNA bands |  |  |
| --- | --- | --- | --- | --- |
|  |  | 40 ng/μl | 20 ng/μl | 10 ng/μl |
| OPW-01 | CTCAGTGTCC | 6 | 6 | 6 |
| OPW-02 | ACCCCGCCAA | 6 | 8 | 8 |
| OPW-03 | GTCCGGAGTG | 4 | 8 | 7 |
| OPW-04 | CAGAAGCGGA | 7 | 9 | 7 |
| OPW-05 | GGCGGATAAG | 4 | 4 | - |
| OPW-06 | AGGCCCGATG | 7 | 8 | - |
| OPW-07 | CTGGACGTCA | 7 | 8 | - |
| OPW-08 | GACTGCCTCT | 5 | 5 | - |
| OPW-09 | GTGACCGAGT | 6 | 6 | - |

The concentration of 20 ng/μl generated the major count of fragments, followed by a concentration of 40 ng/μl. At the concentration of 10 ng/μl, the DNA samples did not amplify with the primers OPW05, OPW06, OPW07, OPW08, and OPW09. Therefore, this is the less indicated of the tested concentrations to be used in the PCR-RAPD reaction with this set of primers and specie, according to our findings.

## 4. Discussion

The relevance to describe an efficient method for DNA extraction for the species *H. commersoni* is justified by the fact that obtaining high-quality samples is crucial for studying organisms at the genetic level. Specifically, in the case of *H. commersoni*, there are few molecular data available in the literature. The information compilated at this work may allow future analyzes of the genetic characterization of the group and, nevertheless, the techniques described here may as well be used to study other fish species.

In this study, we proposed a methodology based on the DNA extraction obtained from ocular tissue, and, to our knowledge, there is no record of this parameter for any fish species to date. The use of this tissue may present some limitations, such as its availability in number and size, but our results indicate that this organ has excellent potential as a source for extract high-quality genomic DNA of *H. commersoni* and possibly for other species.

This technique can be useful in cases where DNA extraction is performed using samples belonging to scientific collections, where the specimens have already been euthanized and are stored for long periods. Potentially, it can also be applied when an alternative model of extraction source is required, once the classical ones (generally from gills, fins, and muscle) have been exhausted without having presented a positive result.

During the test period, all the steps that were shown to be promising regarding the primary objective of our study were selected, aiming to group them into a single protocol, considered the “ideal” one for the *H. commersoni*. In this sense, is possible to discuss some of the characteristics that presented better efficiency in terms of isolated DNA.

The use of liquid nitrogen for tissue maceration in DNA extractions, especially in solid tissues, is a method commonly described as efficient (Chen and Leibenguth, 1995). However, in this work, it was not a factor of influence both on the quality or quantity of product obtained. The non-use of such reagent implies a protocol of lower cost and greater ease of execution, facilitating the application of this methodology within a larger number of laboratories and research groups.

The addition of RNAse is suggested as necessary in protocols for extracting fish DNA, being used to eliminate the contamination of the samples by RNA, consequently increasing DNA purity (Wasko et al., 2003). However, in this study, the use of RNAse did not appear to affect the DNA extraction. For all protocols and variables tested, RNAse did not evolve purer samples. The possibility of eliminating the use of RNAse during DNA extraction also results in the development of a lower cost protocol.

Other variables, however, corroborated with the information available in the literature, such as the use of proteinase K to increase extraction efficiency, especially at temperatures of sample incubation up to 55 ºC. The proteinase K digestion is characterized by the purification of quality DNA for genetic analyzes such as PCR (Sweeney and Walker, 1993; Goldenberger et al., 1995).

One of the most promising results was the possibility of exploring different incubation times of the samples in the lysis buffer, with a proven spectrum of efficiency in one, 12, 24, and 36 hours, using the ocular tissue of *H. commersoni*. The advantages related to this discovery are the possibility of extracting the DNA in a single day (one hour at 55ºC), incubating the samples overnight, or, furthermore, being able to postpone extraction after tissues incubation in lysis buffer, until 36 hours at room temperature of approximately 27 °C. This result implies that the samples can be transported in situations where the specimens are euthanized in the field and the tissue maintained in lysis buffer for up to 36 hours without the need for refrigeration or any other form of preservation until the DNA can be extracted at adequate conditions.

In general, all protocols chosen from literature presented relative efficiency in the extraction of *H. commersoni* DNA, despite some limitations, when evaluating the concentration indices obtained from each sample. However, when the values were analyzed under the spectrum of the absorbance ratio equation, it is evident that the protocol proposed in this work was the one that obtained the highest DNA purity values.

Our improved protocol consists of a simple and economical method to obtain DNA from *H. commersoni*, with the potential to be applied in other similar species. Following this methodology that we optimized, it was possible to amplify the samples using PCR-RAPD primers successfully. The samples from the other protocols were also submitted to the same analysis, but without results. Further studies analyzing the samples obtained using our protocol on different genetic studies, and applying other molecular markers should be conducted in order to expand this discussion.

Under optimal conditions, a single cell contains sufficient DNA to serve as a template for PCR reactions. The efficacy of this protocol in PCR-based genetic studies was confirmed through the amplification of samples using RAPD markers. Our findings brought unprecedented data for the specie *Hypostomus commersoni*, and this information can be applied in further studies of the genetic characterization of this important taxon.

## Acknowledgments

The authors thank CAPES, CNPq, FAPERGS, and the Post-Graduation Program in Ecology from URI (Universidade Regional Integrada-Erechim, Brazil), for the financial support and scholarships provided.

