## Supplementary material for "Evaluation of eight protocols for genomic DNA extraction of *Hypostomus commersoni* Valenciennes, 1836 (Loricariidae: Siluriformes)": Protocols_S1 (from another version of this preprint: version 2, 2020-04-13)

**S1 – Full description of protocols 1 to 7.**

**Protocol 1** **(Doyle and Doyle, 1987):**

a) Reagents: CTAB (2% CTAB + 1,4 M NaCl + 20 mM EDTA + 100 mM Tris-HCl + 1% PVP); proteinase K (1 mg/mL); CIA (chloroform + isoamylic alcohol, 1:24, v/v); TE buffer (10mM Tris + 1 mM EDTA 1 mM pH 8,0); RNAse (1 mg/mL).

b) Steps: 1. Collect 20 mg of sample; 2. Add 500 µL of CTAB, 15 µL of β-Mercaptoetanol and 15 µL of proteinase K and homogenize; 3. Incube at 60 ºC for 1h (or at 50 ºC overnight); 4. Add 500 µL of CIA and homogenize; 5. Centrifuge for 20 minutes at 14.000 rpm; 6. Transfer the aqueous phase to a new tube; 7. Add 400 µL of isopropanol (cold) and homogenize; 8. Storage at -20 ºC for 2 hours; 9. Centrifuge for 20 minutes at 14.000 rpm and discard the supernatant; 10. Wash with 400 µl of ethanol analytical grade; 11. Add 200 µL of absolute ethanol and discard; 12. Dry the *pellet*; 13. Add 50-150 µL of TE and 2 µL of RNAse; 14. Incube for 1 h (37 ºC) and 24 hours at 25-27 ºC.

**Protocol 2** **(Medrano et al., 1990):**

a) Reagents: STE buffer (0,1 M NaCl + 10mM Tris + 1 mM EDTA pH 8,0); lysis buffer (0,1 M NaCl + 50 mM Tris + 50 mM EDTA pH 8,0); 20% SDS (2 g/mL); proteinase K (1 mg/mL); 4M NaCl (11,71 g/50 mL); TE buffer (10 mM Tris + 1 mM EDTA pH 8,0); RNAse (1 mg/mL).

b) Steps: 1. Collect 20 mg of sample; 2. Wash with 500 µL of STE buffer; 3. Add 550 µL of lysis buffer, 27,5 µL of 20% SDS and 40 µL of proteinase K; 4. Incube at 60 ºC for 2 hours (vortex each 15 minutes); 5. Add 600 µL of 4M NaCl; 6. Mix in vortex for 15 seconds and centrifuge for 30 minutes at 13.000 rpm; 7. Transfer the supernatant to a new tube; 8. Add 400 µL of isopropanol (cold) and homogenize; 9. Incube overnight at 4 ºC or for 2 hours at -20 ºC; 10. Centrifuge for 30 minutes at 14.000 rpm and discard the supernatant; 11. Add 700 µL of 70% ethanol and homogenize; 12. Centrifuge for 5 minutes at 9.000 rpm; 13. Discard and incube the tubes at 37 ºC for 30 minutes; 14. Leave the samples 10 minutes on the bench; 15. Add 30 µL TE buffer; 16. Incube for 1 hora at 37 ºC; 17. Dry the pellet for 12 hours at 25-27 ºC; 17. Storage at 4 ºC.

**Protocol 3** **(Whitmore et al., 1992):**

a) Reagents: lysis buffer (100 mM Tris-HCl pH 8,0 + 10 mM EDTA + 0,1% SDS + 50 mM DTT); proteinase K (0,5 mg/mL); phenol (saturated with Tris, pH 8,0); CIA (chloroform + isoamylic alcohol, 1:24 v/v).

b) Steps: 1. Collect 20 mg of sample; 2. Add 500 µL of lysis buffer and 14 µL of proteinase K; 3. Incube for 4 hours at 50 ºC or overnight at 42 ºC; 4. Add 700 µL of phenol, centrifuge for 15 minutes at 9.000 rpm and transfer the supernatant to a new tube; 5. Add 700 µL of CIA and centrifuge for 15 minutes at 6.000 rpm; 6. Transfer the supernatant to a new tube; 7. Add 600 µL absolute ethanol; 8. Centrifuge for 10 minutes at 7.000 rpm; 8. Discard the ethanol and dry the tube on the bench; 9. Add 200 µL of distilled water to the pellet; 10. Incube at 25-27 ºC for 12 hours; 11. Storage at -20 ºC.

**Protocol 4 (****Aljanabi and Martinez, 1997):**

a) Reagents: lysis buffer (0,4 M NaCl + 10 mM Tris-HCl pH 8,0 + 2 mM EDTA pH 8,0); 20% SDS (2 g/mL); proteinase K (1 mg/mL); 4M NaCl (11,71 g/50 mL);

b) Steps: 1. Collect 20 mg of sample; 2. Add 400 µL of lysis buffer and homogenize in vortex for 15 seconds; 3. Add 400 µL of 20% SDS and 80 µL of proteinase K; 4. Incube at 35 ºC for 1 hora or overnight at 55–65 °C; 5. Ad 300 µL of 4 M NaCl and vortex; 6. Transfer the supernatant to new tubes. 7. Add 600 µL of isopropanol and homogenize; 8. Incube for 1 hour at –20 ºC; 9. Centrifuge for 20 min at 11.000 rpm; 10. Discard the supernatant, add 400 µL of ethanol analytical grade and discard; 11. Dry at 25-27 ºC; 12. Add 300-500 µL of distilled sterile water; 13. Storage at 4 ºC.

**Protocol 5 (****Faleiro et al., 2003):**

a) Reagents: lysis buffer (100 mM Tris-HC1 pH 8,0 + 20 mM EDTA + 2,8% CTAB + 1,3 M NaC1 + 1% PVP + 0,2% β-Mercaptoetanol); CIA (chloroform + isoamylic alcohol 1:24 v/v); 7% CTAB (7% CTAB + 1,4 M NaCl + 20 mM EDTA + 100 mM Tris-HCl + 1% PVP); TE buffer (10 mM Tris pH 8,0 + 1 mM EDTA).

b) Steps: 1. Collet 20 mg of sample and macerate with liquid nitrogen; 2. Transfer to a tube; 3. Add 800 µL of lysis buffer, homogenize; 4. Incube for 1 hour at 70 ºC; 5. Ad 700 µL of CIA and mixed for 10 minutes; 6. Centrifuge for 15 minutes at 13.000 rpm. 7. Transfer the supernatant and add 55 µL of 7% CTAB; 8. Repeat the steps 5, 6 and 7; 9. Add 700 µL of absolute isopropanol.; 10. Incube for 2 hours at -20 ºC and centrifuge for 15 minutes at 13.000 rpm; 11. Discard the supernatant and wash the pellet in 700 µL of ethanol analytical grade (repeat); 13. Discard the volume and dry the pellet; 14. Add 100 µL of TE buffer; 15. Storage.

**Protocol 6** **(Barrero et al., 2008):**

a) Reagents: lysis buffer (50 mM Tris-HC1 pH 8,0 + 50 mM EDTA + 100 mM NaC1); 1% SDS (1 g/100 ml); 4M NaCl (11,71 g/ 50 mL); proteinase K (20 mg/mL); TE buffer (10 mM Tris pH 8,0 + 1 mM EDTA); RNAse (30 µg/mL);

b) Steps: 1. Collect the samples, add 550 μl of lysis buffer, 7 µl of 1% SDS and 20 µl of proteinase K; 2. Incube for 12 hours at 50 °C; 3. Add 600 μl of 5 M NaCl and centrifuge for 10 minutes at 12.000 rpm; 4. Transfer the supernatant and add 700 μl of ethanol analytical grade; 5. Incube for 2 hours at -20 °C; 6. Centrifuge for 20 minutes at 12.000 rpm; 7. Add 700 μl of ethanol analytical grade and discard; 8. Dry the pellet; 9. Add 80 µL of TE buffer; 10. Add 3 μl of RNAse; 11. Incube for 40 min at 37 ºC; 12. Storage at -20 ºC.

**Protocol 7 (****Mossi et al., 2014):**

a) Reagents: STE buffer (0,1 M NaCl + 10 mM Tris + 1 mM EDTA pH 8,0); lysis buffer (1 M Tris-Hcl pH 8,0 + 0,5 M EDTA pH 8,0 + 10% SDS + 5 M NaCl + 1% β-Mercaptoetanol); proteinase K (1 mg/mL); CIA (chloroform + isoamylic alcohol, 1:24 v/v); TE buffer (1 M Tris pH 8,0 + 0,5 M EDTA pH 8,0).

b) Steps: 1. Collet 20 mg of sample; 2. Add 1 mL of STE (cold) and remove it, repeating this step 3x; 3. Add 500 µL of lysis buffer and 11 µL of proteinase K; 4. Vortex and incubate overnight at 37 ºC or 30 minutes at 55 ºC; 5. Add 500 µL of CIA, homogenize by inversion for 45 seconds and centrifuge for 10 minutes at 14.000 rpm; 6. Transfer 450 µL of supernatant, add 900 µL of isopropanol and homogenize. 7. Incube for 2 hours at 20 ºC or overnight at 4 ºC; 8. Centrifuge for 15 minutes at 14.000 rpm. Discard the supernatant and dry the pellet; 9. Add 1 mL of ethanol analytical grade and centrifuge for 5 minutes at 6.000 rpm; 10. Discard the supernatant and dry the pellet again; 11. Add 100 µL of TE and incube for 2 hours at 37 ºC; 12. Storage at 4 ºC.
